# Uncovering long-term existence of a silent short-term memory trace

**DOI:** 10.1101/2021.05.08.443276

**Authors:** Maha E. Wally, Masanori Nomoto, Kareem Abdou, Kaoru Inokuchi

**Affiliations:** Research Center for Idling Brain Science, University of Toyama, Toyama 930-0194, Japan; Department of Biochemistry, Graduate School of Medicine and Pharmaceutical Sciences, University of Toyama, Toyama 930-0194, Japan; CREST, JST, University of Toyama, Toyama 930-0194, Japan; Pharmacology Department, Faculty of Pharmacy, The British University in Egypt, Cairo 11837, Egypt; Department of Biochemistry, Faculty of Pharmacy, Cairo University, Cairo 11562, Egypt

## Abstract

Active recall of short-term memory (STM) is known to last for a few hours, but whether STM has long-term functions is unknown. Here we show that, STM can be optogenetically retrieved at a time point during which natural recall is not possible, uncovering the long-term existence of a silent STM engram. Moreover, re-training within 3 days led to natural long-term recall, indicating facilitated consolidation. Calcium imaging revealed hippocampal CA1 reactivations of the STM trace during post-learning sleep. Inhibiting offline CA1 activity, N-methyl-D-aspartate receptor activity, or protein synthesis after first exposure to the STM-forming event impaired the future re-exposure-facilitated consolidation, which highlights a role of protein synthesis and sleep in storing a silent STM trace. These results provide evidence that STM is not completely lost within hours and demonstrates a possible two-step STM consolidation, first storage as a silent engram, then transformation into an active state by recurrence within 3 days.

## Introduction

Short-term memory (STM) is formed by the delivery of a weak stimulus while long-term memory (LTM) is formed by a stronger and more enduring stimuli^1^. Moreover, STM is known to last for only a few hours, whereas LTM is subjected to a consolidation process that allows it to remain for long periods of time^2–5^. However, it is not yet clear whether STM has long-term effects beyond this time.

Engram cells are neuron populations in which memories are stored, and their specific reactivation leads to individual memory retrieval^6,7^. Recent advances and new genetic technologies enabled the use of immediate early genes^8^, such as *c-fos*, as markers of neuronal activity to visualize and specifically label the cellular ensembles constituting the engram^9^. This gave rise to a wide range of studies that manipulated memory engrams through erasure^10^, stimulation^11^ or even creation of false memories^12,13^. Engram cells were also shown to be involved in memory allocation and memory association^14–16^. Furthermore, engram studies enabled the identification of the synaptic correlate of specific memories^17,18^. Two major types of engrams have been defined: the active engram, which can be naturally recalled, and the silent engram, which cannot be retrieved by natural cues, but can be activated by artificial stimulation^19,20^. Silent engrams have been shown to be present in cases of retrograde amnesia^21,22^ and early-onset Alzheimer’s disease^23^. Moreover, silent engrams are involved in the reorganization of remote memories between the hippocampus and cortex through system consolidation^19,24^. However, all engram-based studies have focused on several types of LTM, and none have investigated the existence of such populations in case of STM.

Memory consolidation process depends on new protein synthesis, whereby the inhibition of post-learning protein synthesis blocks LTM but not STM retrieval^25,26^. LTM consolidation has also been shown to occur after a period of post-learning sleep^27^ which consists of two main stages, non-rapid eye movement sleep (NREM) characterized by slow delta waves, spindles and sharp wave-ripples (SWRs), and rapid-eye sleep (REM) characterized by fast theta rhythms^28^. It has been shown that hippocampal SWRs, which occur during NREM sleep, and theta rhythms, which occur during REM sleep, are both critical for the consolidation of spatial memories^29,30^. Furthermore, memory reactivation has been suggested as one of the mechanisms underlying memory consolidation during post-learning sleep^31–33^ and awake states^34,35^.

We hypothesized that, beyond the few hours of active STM recall, a trace of STM may continue to exist in a silent manner, unable to be naturally retrieved, but its activation could have long-term effects, nevertheless. We investigated this hypothesis using optogenetic recall of an STM-forming event, which unveiled the presence of a silent STM engram stored within the hippocampal circuit. Using a simple behavioral approach, we demonstrate that the silent STM engram provides a template that facilitates consolidation upon future re-exposure to the same event within a specific period of time. Using pharmacological manipulations, we investigated the mechanisms behind the long-term storage of the silent STM trace. Contrary to the previously accepted concept, STM triggers new protein synthesis for the storage of its silent engram. N-methyl-D-aspartate receptor (NMDAR) is involved in facilitating STM consolidation. Furthermore, using *in vivo* calcium imaging we show reactivation of the STM trace during post-learning sessions including sleep. Finally, we highlight a necessity of post-learning sleep in preserving the availability of the silent STM trace in a similar fashion to its consolidation of LTM trace using specific optogenetic silencing of offline activity during post-learning NREM sleep. Taken together, these results modify our current understanding of the STM and provide mechanistic insights into its silent storage and potential long-term effects.

## Results

### LTM and STM in novel object location task

We used the novel object location (NOL) behavioral paradigm^23^, a hippocampal-dependent spatial memory^36^, as a learning task. Mice freely explored the location of two objects during a training session, and then during the test session one object was moved to a new location while the other remained in its original location (Extended Data Fig. 1). When mice recall the memory of object location, their exploration of the moved object is greater than that of the unmoved object. However, an equal exploration of the two objects indicates that the mice have not discriminated between the new and old locations, and have been unable to retrieve the memory. We used two training protocols^37^, as follows: a strong protocol for 15 min (Fig. 1a) and a weak protocol for 6 min (Fig. 1d). The strong training produced a LTM that was successfully recalled after 1 day (Fig. 1c), and the weak training produced a STM that was recalled within 30 min, but not after 1 day (Fig. 1f). Lack of initial bias to one of the two objects during the training sessions was confirmed for all groups (Fig. 1b,e).

**Fig. 1.**
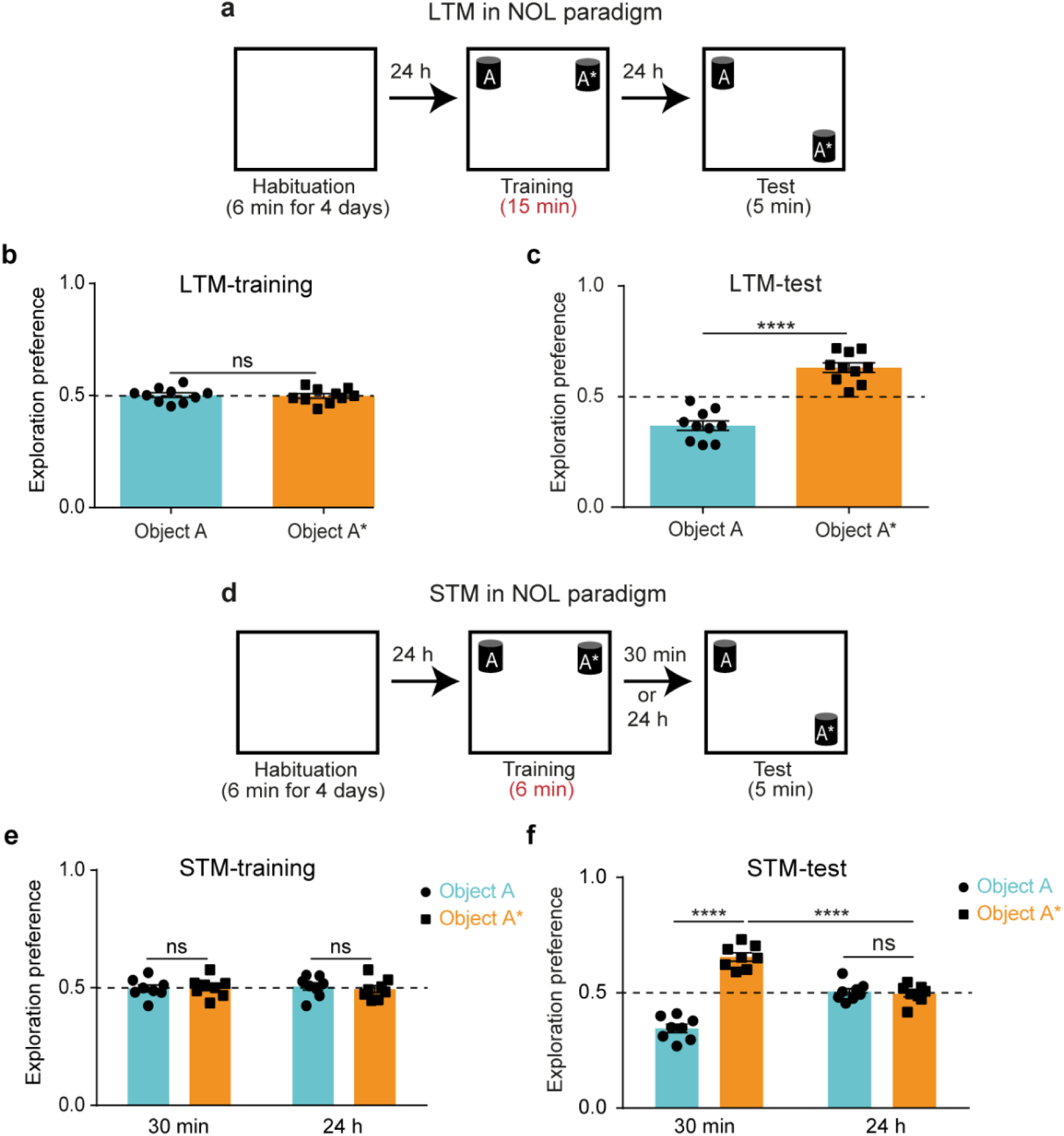
LTM and STM in novel object location task (NOL). **a,** NOL paradigm with the strong training protocol (15 minutes). **b and c,** Exploration preference for each object during the strong training **(b)** and the test sessions **(c)** (n = 10). **d,** NOL paradigm with the weak training protocol (6 minutes). **e and f,** Exploration preference for each object during the weak training **(e)** and the test sessions **(f)** for groups tested after 30 minutes (n = 8) or 24 hours (n = 8). Comparisons were made using unpaired student’s t-test; ns, not significant (*P* > 0.05), \*\*\*\**P* < 0.0001. Data are presented as the mean ± SEM.

### Optogenetic recall of the silent STM engram

Using the weak training protocol, we investigated whether STM has an engram by performing optogenetic activation of the STM 1 day after its initial acquisition, a time point at which mice do not retrieve the memory (Fig. 1f). We used c-fos::tTA/KA1::Cre double transgenic mice injected with AAV_9_-TRE-DIO-ChR2-mCherry into their hippocampal CA3 to specifically label activated CA3 cells involved in learning the weak event with the blue-light sensitive ChR2^38^. One day after training, mice were subjected to the test session, during which optical stimulation to their CA3-CA1 projections was conducted by shining blue light (473 nm) above the CA1 area at 20 Hz stimulation (Fig. 2a-c). Mice that received this light stimulation (light ON group) explored the moved object significantly more than the unmoved object, compared to the control group (light OFF group) (Fig. 2d), which indicates the successful long-term recall of their STM. This result indicates that similar to the previously reported cases of retrograde amnesia and early-onset Alzheimer’s disease^21,23^, STM also forms a silent engram which can be artificially recalled 1 day later and that this engram is stored within the hippocampal CA3-CA1 circuit.

**Fig. 2.**
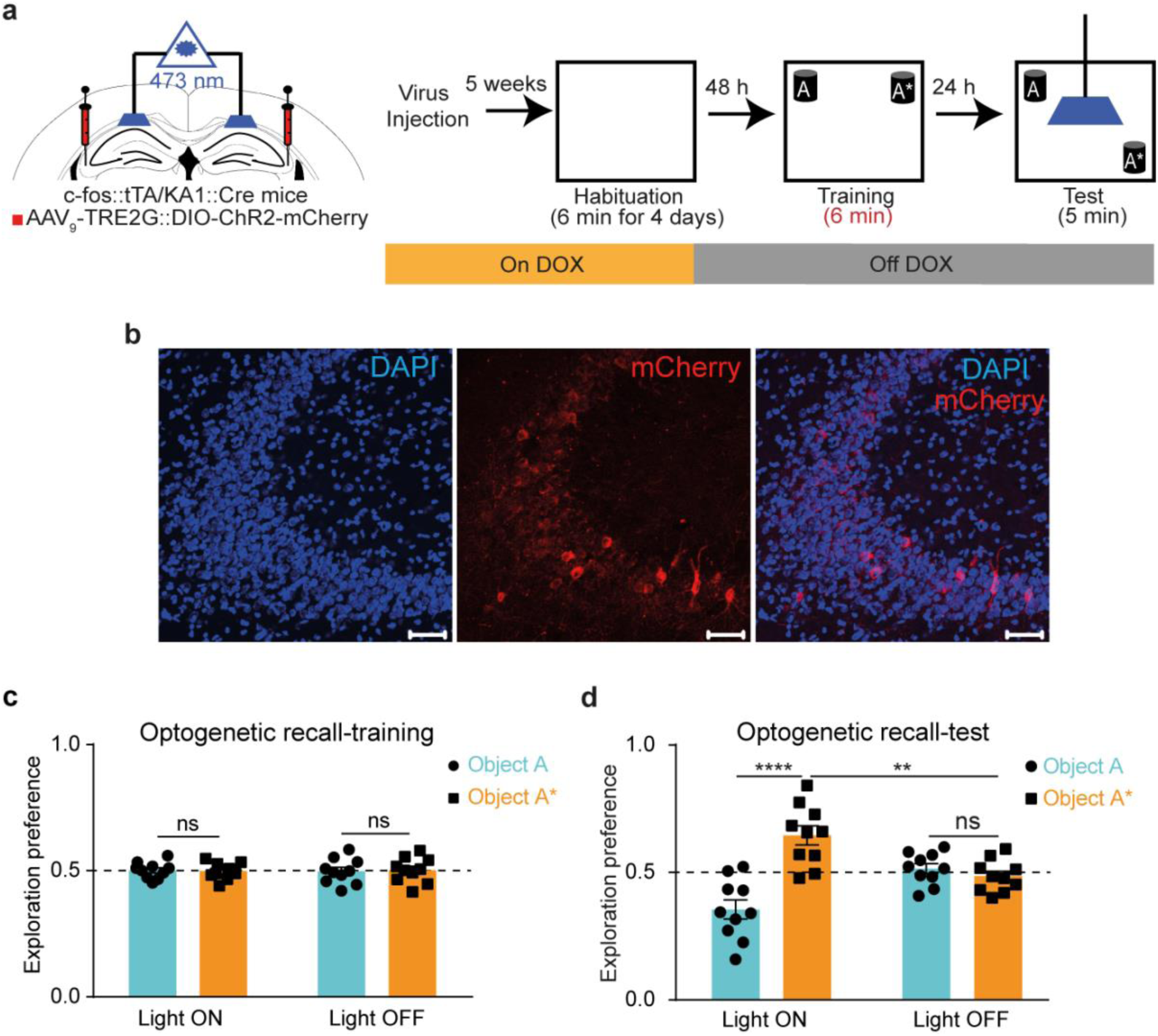
Optogenetic recall of the silent STM engram. **a,** Left: Diagram of AAV injection into CA3, guide cannula and optic fiber placement into CA1. Right: Experimental design for labelling and optogenetic recall of the NOL STM engram. **b,** Representative section of CA3 showing mCherry protein expression (scale bar = 50 μm, DAPI: 4′,6-diamidino-2-phenylindole). **c and d,** Exploration preference for each object during the training **(c)** and the test **(d)** sessions in light ON (n = 10) vs. light OFF (n = 10) groups. Comparisons were made using unpaired student’s t-test; ns, not significant (*P* > 0.05), \*\**P* < 0.01, \*\*\*\**P* < 0.0001. Data are presented as the mean ± SEM.

### Re-exposure facilitates STM consolidation

To test whether the silent STM engram can have long-term effects we tested the possibility that this engram can become activated by re-exposure to the same event in the future and whether this activation can lead to consolidation and natural long-term recall. Mice underwent the weak NOL training, and then, 1 day later, repeated the same training again and were then tested for consolidation after one more day (Fig. 3a-c). Mice subjected to this two-training paradigm succeeded to recall the STM 1 day after re-exposure, which indicates consolidation of the STM trace (Fig. 3d). This result demonstrates that the silent STM engram, produced by the first weak training, was used as a template by the second weak training to facilitate consolidation. Without the presence of the initial STM engram, the second weak event would have been processed independently and would have produced a STM, unable to be naturally retrieved one day later.

**Fig. 3.**
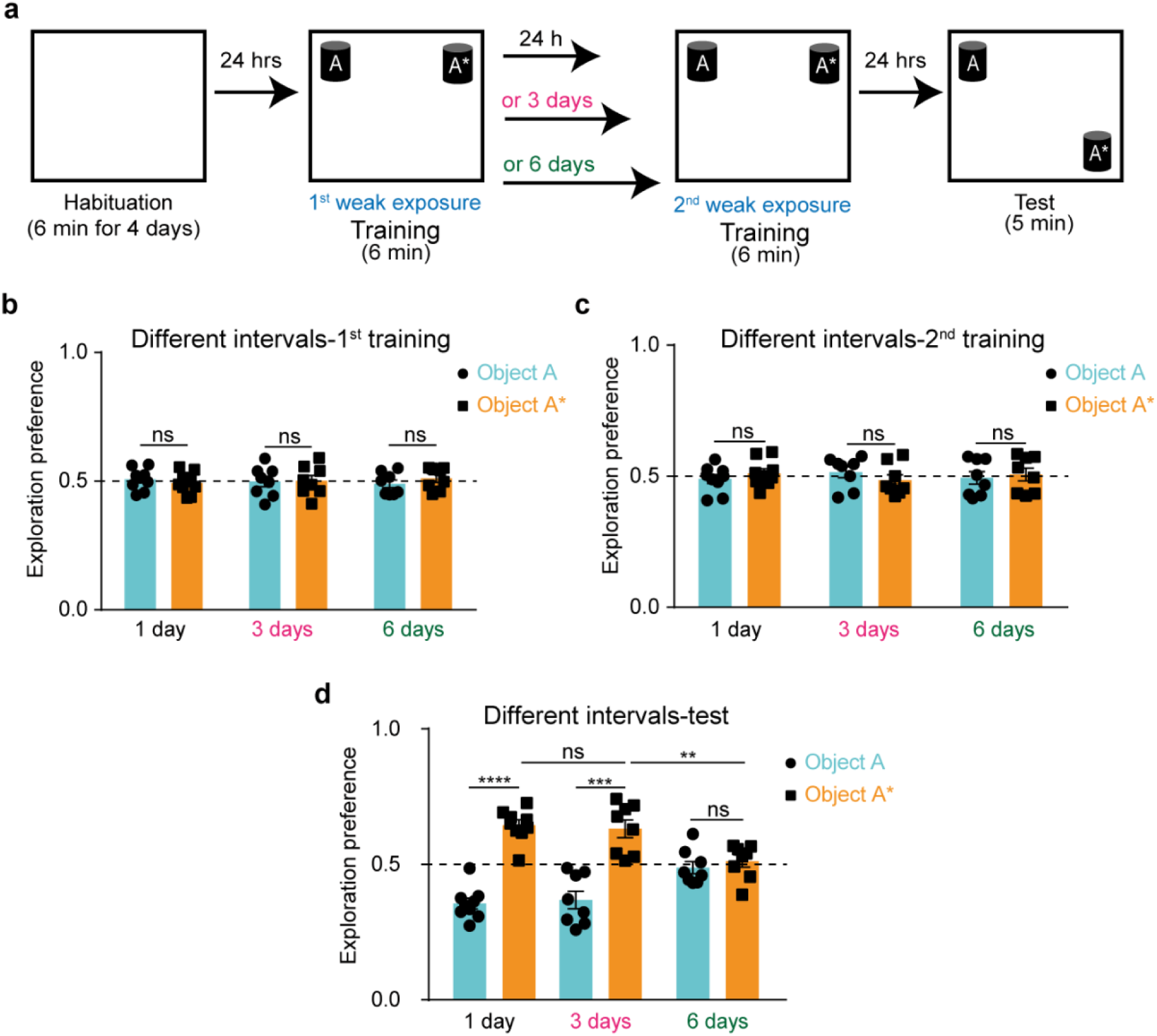
Re-exposure facilitates STM consolidation. **a,** NOL two-training paradigm with different intervals in between. **b, c and d,** Exploration preference for each object during the first training **(b)**, the second training **(c)** and the test **(d)** sessions for different intervals groups; 1-day (n = 9), 3-day (n = 8), and 6-day (n = 8). Comparisons were made using two-way analysis of variance (ANOVA) followed by Bonferroni Post-hoc test; ns, not significant (*P* > 0.05), \*\**P* < 0.01, \*\*\**P* < 0.001, \*\*\*\**P* < 0.0001.

To assess how long the silent STM engram remains available for this future re-usage, we implemented the two-training paradigm while increasing the spacing interval between the two training sessions (Fig. 3a-c). Using a 3-day interval, mice still recalled the memory similar to the 1-day interval; however, with a 6-day interval, the memory retrieval was no longer observed (Fig. 3d). These results indicate that the silent STM engram remains available for activation and subsequent consolidation if the same experience is repeated within 3 days.

### Storage of the silent STM trace is protein-synthesis dependent

One of the major distinctions between STM and LTM is that LTM storage depends on new protein synthesis, but STM does not^25,26^. These new proteins are believed to alter the shape and function of the involved synapses which contributes to the long retention time of LTM^5^. However, in case of STM, it is believed that new protein synthesis does not occur, but rather the covalent modification of pre-existing proteins causes temporary amplification of the synaptic transmission that contributes to the short retention time of STM^39^.

Our finding that STM forms an engram made us curious as to whether this engram requires new proteins for its storage. We repeated the two-training paradigm with a 1-day interval and injected anisomycin, a protein synthesis inhibitor, to the hippocampal CA1 region immediately after the first training session (Fig. 4a-c and Extended Data Fig. 2). Mice injected with anisomycin did not recall the STM, unlike the PBS control group (Fig. 4d), which suggests that anisomycin disrupted the storage of the silent STM engram produced by the initial training, which led to failure in its consolidation after recurrence. This shows that while new protein synthesis is not involved in the same-day recall of STM, it is required for the long-term storage of its silent trace.

**Fig. 4.**
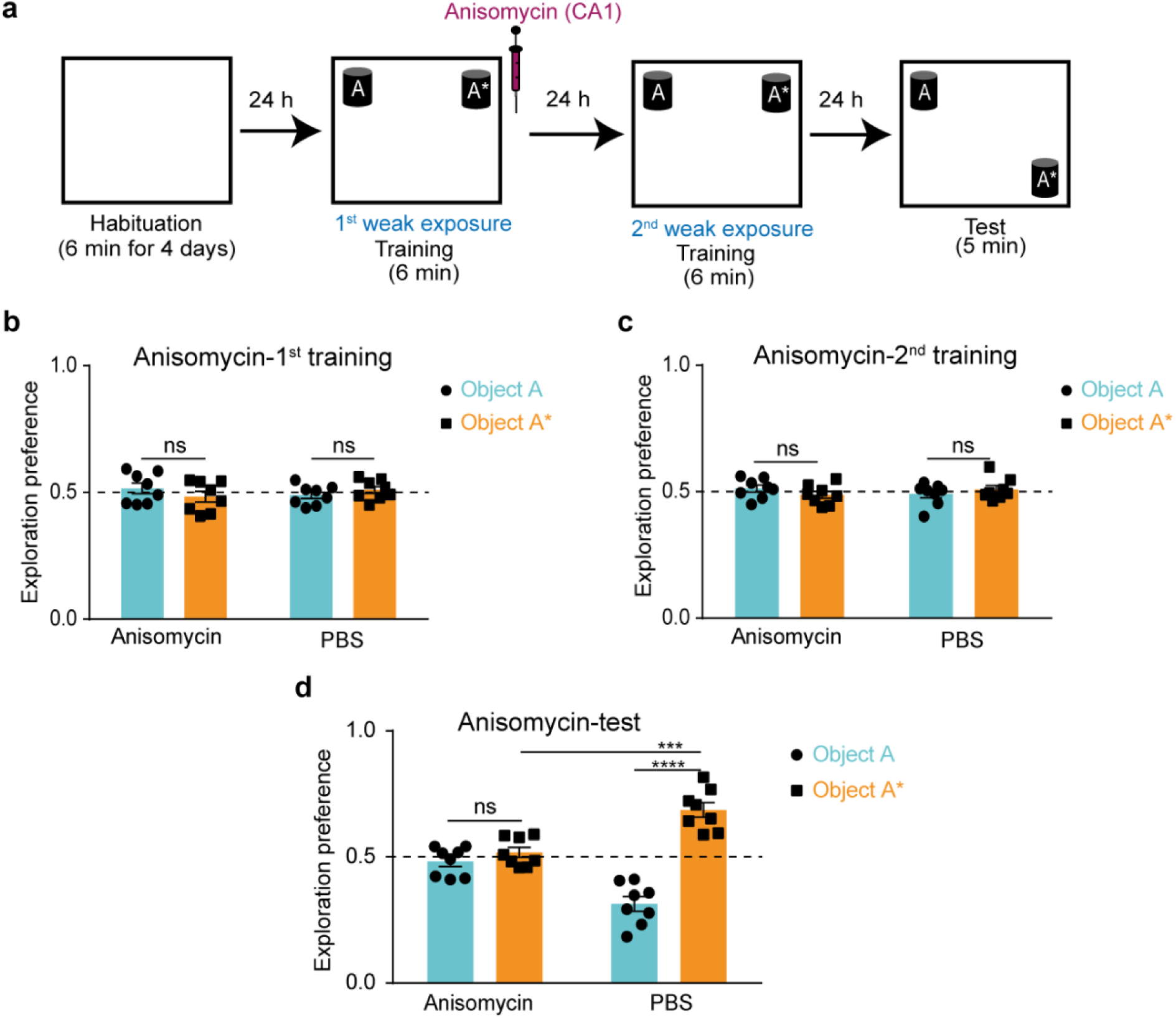
Storage of the silent STM trace is protein-synthesis dependent. **a,** NOL two-training paradigm with 1-day interval in between with anisomycin injection after the first training session into the CA1 region. **b and c,** Exploration preference for each object during the first **(b)** and the second **(c)** training sessions for anisomycin- (n = 8) and PBS- (n = 8) injected groups. **d,** Exploration preference for each object during the test session for anisomycin- (n = 8) and PBS- (n = 8) injected groups. Comparisons were made using unpaired student’s t-test; ns, not significant (*P* > 0.05), \*\*\**P* < 0.001, \*\*\*\**P* < 0.0001. Data are presented as the mean ± SEM.

### Storage and activation of the silent STM trace is N-methyl-D-aspartate receptor-dependent

NMDAR, the ionotropic glutamate receptor, plays a major role in memory processing where its activation leads to Ca^2+^/calmodulin-dependent protein kinase II (CaMKII) phosphorylation, which in turn triggers a cascade of molecular events through which long-term potentiation (LTP) and synaptic plasticity occurs in the dendritic spines^5,40,41^; the storage site of specific memories^17^. D-AP5 is an NMDAR blocker that prevents the LTP of synaptic transmission and has been used to examine the role of NMDAR in learning and memory^42,43^. Similar to the anisomycin experiment, using the two-training paradigm with a 1-day interval, mice injected with AP5, in the hippocampal CA1 region after either the first or second weak trainings (Fig. 5a and Extended Data Figs. 3, 4, and 5) did not differentiate between the two objects in the test session compared with the PBS control groups (Fig. 5b,c), which indicates failure in the storage and activation of the silent STM engram, respectively. This further indicates that the NMDAR signaling pathway, and potentially synaptic plasticity, are involved in the facilitated consolidation of the STM by storing and activating the silent STM trace, consecutively.

**Fig. 5.**
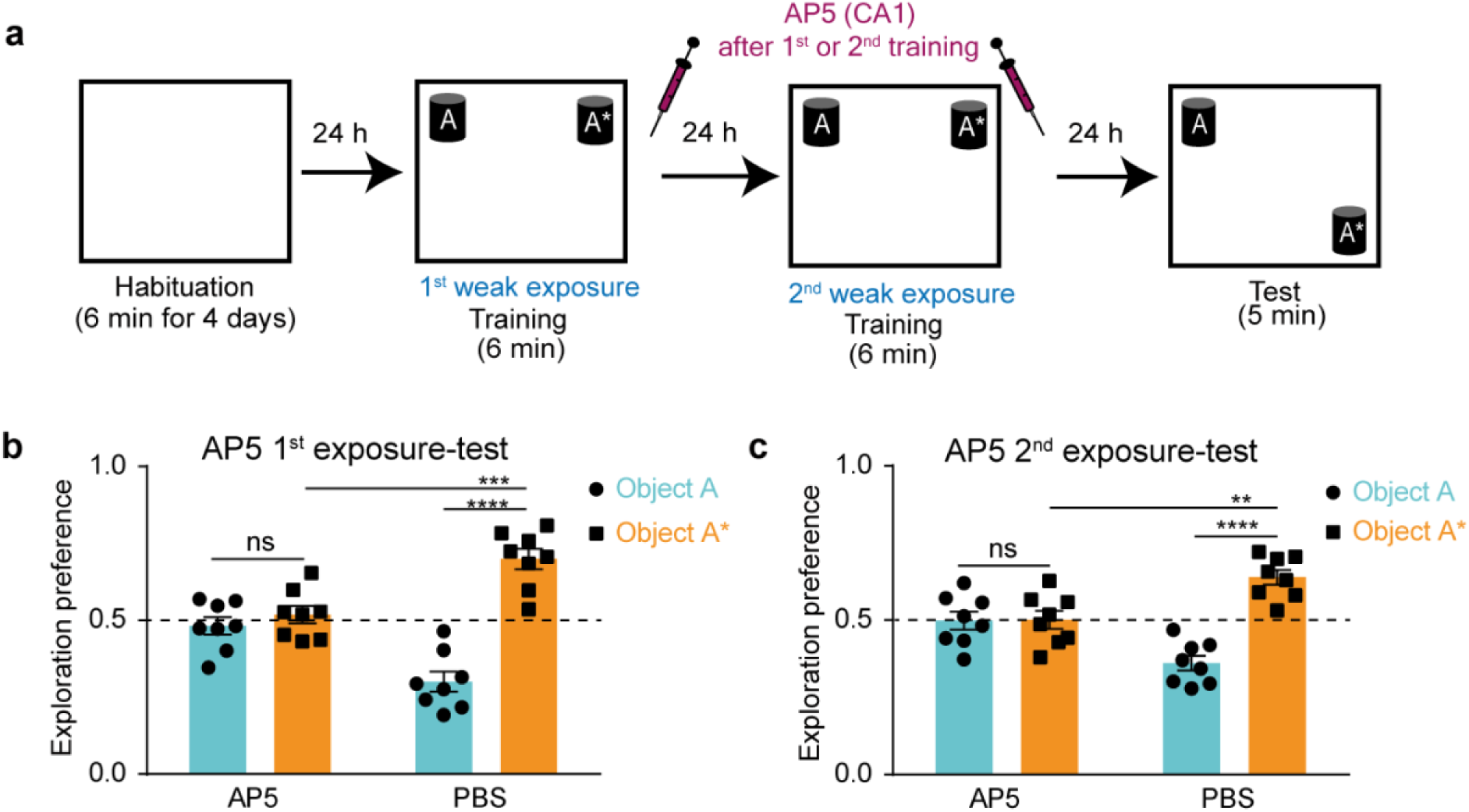
Storage and activation of the silent STM trace is N-methyl-D-aspartate receptor-dependent. **a,** NOL two-training paradigm with 1-day interval in between with AP5 injection after either the first or the second training sessions into the CA1 region. **b and c,** Exploration preference for each object during the test session for groups injected after the first **(b)** or the second **(c)** training sessions for AP5- (n = 8) and PBS- (n = 8) injected groups. Comparisons were made using unpaired student’s t-test; ns, not significant (*P* > 0.05), \*\**P* < 0.01, \*\*\**P* < 0.001\*\*\*\**P* < 0.0001. Data are presented as the mean ± SEM.

### STM trace reactivates during post-learning NREM sleep

To further investigate the underlying mechanism of the silent STM trace storage, *in vivo* calcium imaging was employed. Recording of the same hippocampal CA1 neurons^44^ was conducted during the two-training NOL paradigm with a 1-day interval and during the post-learning NREM sleep sessions as well (Fig. 6a-c; see Methods). An automated sorting system^45^ was used to extract each neuron’s calcium activity and the mean neuronal activity for each session was calculated using mean z-score calculations (see Methods). Accordingly, two population of neurons were identified; the first population showed high calcium transients starting from the habituation session (habituation-emerging neurons), while the second population showed low calcium transients during the habituation session; however, from the first training, calcium activity of this second population significantly increased (training-1-emerging neurons) (Fig. 6d,e; see Methods). The sustained activity of training-1-emerging neurons throughout the following sessions: post-learning NREM sleep-1, training-2, post-learning NREM sleep-2, and test (Fig. 6e), suggested a possible reactivation of STM similar to the previously reported replay of LTM^31^.

**Fig. 6.**
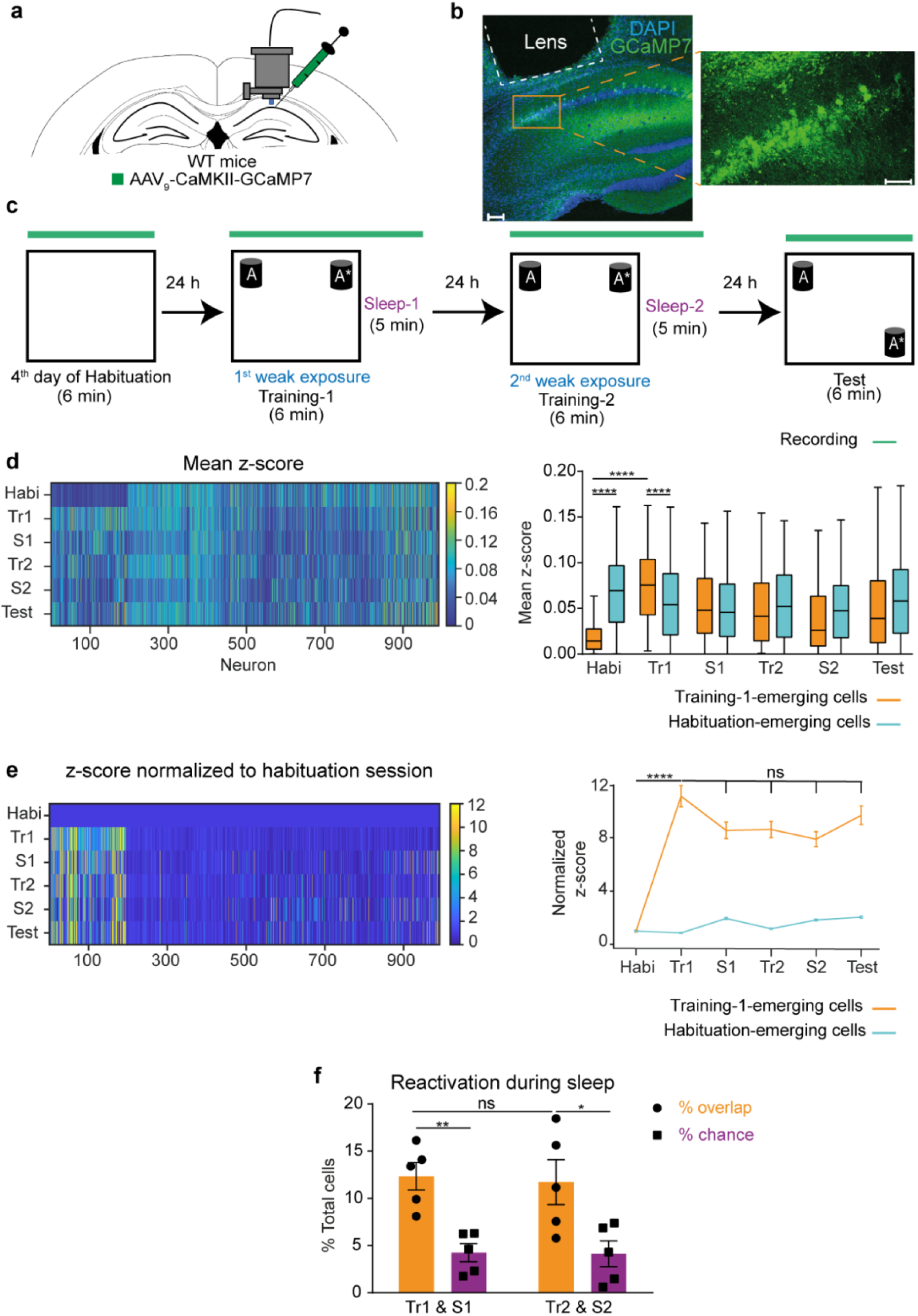
STM trace reactivates during post-learning NREM sleep. **a,** Diagram of AAV injection and miniature microscope placement into CA1. **b,** Representative section of CA1 showing GCaMP7 protein expression (scale bar left = 100 μm, right = 50 μm). **c,** Calcium imaging of NOL two-training paradigm. **d,** Left: mean z-score of neurons (n = 991 > 3 SD from five mice) during different sessions. (Bottom: number of neurons, right: z-score scale, left: imaging sessions; Habi: habituation, Tr1: training-1, S1: NREM sleep-1, Tr2: training-2, S2: NREM sleep-2). Right: mean z-score for training-1-emerging cells (n = 195) vs. habituation-emerging cells (n = 796). Comparisons were made using Wilcoxon rank sum and Wilcoxon matched-pairs signed rank test; \*\*\*\**P* < 0.0001. In box plot, center line represents the median, box edges represent top and bottom quartiles, whiskers represent maximum and minimum values. **e,** Left: mean z-score as in **(d)** normalized to that of the habituation session. Right: mean z-score normalized for training-1-emerging cells vs. habituation-emerging cells. Comparisons were made using paired Student’s t-test; ns, not significant (*P* > 0.05), \*\*\*\**P* < 0.0001. **f,** Percentage of cells overlapping between training-1 session and NREM sleep-1 session (Tr1&S1) vs. training-2 session and NREM sleep-2 session (Tr2&S2) compared with chance (n = 5 mice, see Methods for details). Comparisons were made using unpaired Student’s t-test; ns, not significant (*P* > 0.05), \**P* < 0.05, \*\**P* < 0.01. Data are presented as the mean ± SEM (**(e)** right, **(f)**).

To investigate whether the STM trace is reactivated during post-learning sleep sessions, the percentage of overlapping neurons between the training sessions and their corresponding sleep sessions was calculated and was found to be significantly higher than the chance levels (Fig. 6f; see Methods). These results suggest that a similar mechanism of reactivation underlies the storge and consolidation of the silent STM trace before and after its activation, respectively, which could reflect the necessity of post-learning sleep in preserving the STM trace, as with the LTM trace^28–30^.

### Causal link between offline activity during post-learning NREM sleep and long-term storage of the silent STM trace

To confirm the necessity of post-learning sleep in preserving the silent STM trace, we used the two-training paradigm with a 1-day interval in between, and subjected mice to a 5 h sleep deprivation immediately after the first training (Extended Data Fig. 6a-c). Contrary to the previous result (Fig. 3d), mice did not discriminate between the two objects during the test session (Extended Data Fig. 6d), which demonstrates a necessity for a post-learning sleep period to preserve the silent STM trace.

For a more specific approach, a group of mice was injected with AAV_9_-CaMKII-ArchT-eYFP (test group) and another group was injected with AAV_1_-CaMKII-eYFP (control group). Orange light (589 nm) was delivered to their hippocampal CA1 region during all the NREM sleep stages that appeared within a 2-hour sleep session immediately after the first training to optogenetically silence the offline hippocampal CA1 neurons (Fig. 7a-f; see Methods). In line with the sleep deprivation results, ArchT-expressing mice failed to recall the memory compared with the eYFP-expressing control group (Fig. 7g) even though both groups had comparable duration of NREM sleep (Fig. 7h). Taken together, these results provide a causal link between post-learning offline activity and the long-term storage of a silent STM trace.

**Fig. 7.**
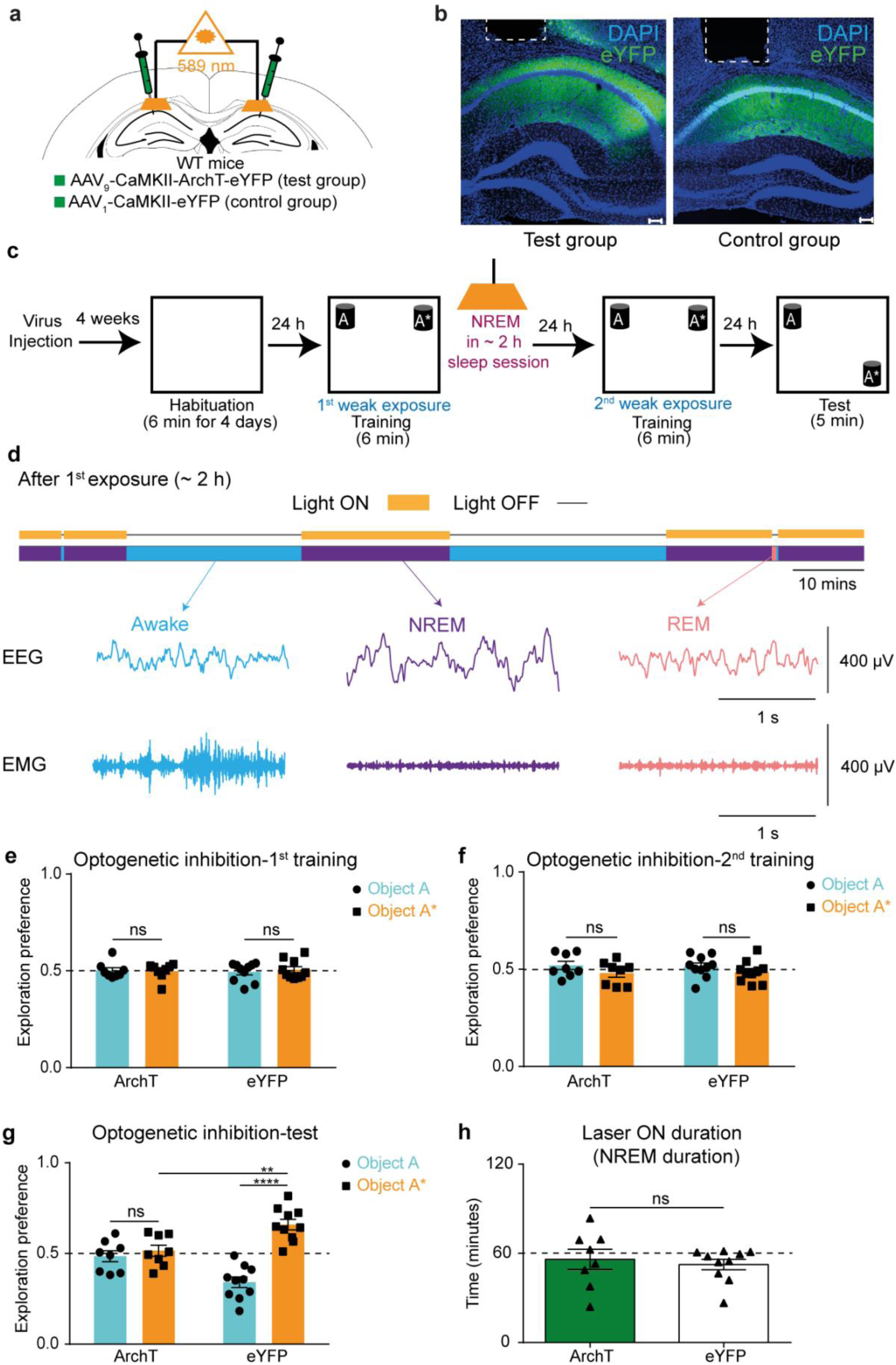
Causal link between offline activity during post-learning NREM sleep and long-term storage of the silent STM trace. **a,** Diagram of AAV injection and CA1 optic fiber placement. **b,** Representative CA1 sections showing ArchT-eYFP (Left: test group) and eYFP (Right: control group) protein expressions (scale bar = 100 μm; dashed line: guide cannula location). **c,** Experimental design for optogenetic silencing of STM trace storage using NOL two-training paradigm during post-learning NREM sleep. **d,** Example from one mouse showing 2-hour EEG/EMG recordings with laser ON during NREM state only (top), and example of EEG/EMG traces for NREM, REM, and awake states (bottom) (see Methods for details). **e and f,** Exploration preference for each object during the first **(e)** and the second **(f)** training sessions for ArchT (n = 8) and eYFP (n = 10) groups. **g** Exploration preference for each object during the test session in ArchT (n = 8) and eYFP (n = 10) groups. **h,** Duration of NREM sleep (laser ON) in ArchT (n = 8) and eYFP (n = 10) groups. Comparisons were made using unpaired Student’s t-test; ns, not significant (*P* > 0.05), \*\**P* < 0.01, \*\*\*\**P* < 0.0001. Data are presented as the mean ± SEM.

## Discussion

Previous studies have extensively researched STM formation, the molecular pathways involved, and its distinctions from LTM^2–5^. However, the nature of the STM engram-wise and the fate of its trace beyond the few hours of its active recall remain unclear. Here, we demonstrate that the STM trace is not entirely erased from the brain within a few hours, but continues to exist in a silent state. Furthermore, we found that the silent STM engram becomes consolidated when the same experience is encountered within 1 to 3 days. This suggests that most likely our brain keeps a record of day-to-day weak events as a silent template that primes the consolidation or strengthening of similar or reoccurring experiences in the near future.

Classical studies have shown that protein synthesis is not needed for the recall of novel weak experiences; however, these studies focused on active STM recall hours from encoding^25,26,46^. We showed that STM does in fact trigger new protein synthesis crucial for the storage and future activation of its silent engram. This result also provides the missing link between our finding that STM forms an engram and the classical notion that it does not trigger new protein synthesis. Thus, our results complement previous work by indicating that a weak event triggers new protein synthesis that is dispensable for active recall of the same memory on the same day, but crucial for the temporary storage of this event in a silent and reversible trace that facilitates future consolidation in the case of reoccurrence. However, future studies are needed to identify the proteins recruited in the storage of the silent STM engram and whether they are different from those recruited in LTM consolidation. Also, a correlation between the duration of the silent engram availability and the half-lives of these proteins remains to be established.

Our finding that NMDAR blockade affects both the storage and activation of the silent STM engram suggests that a two-step process of synaptic plasticity may occur: a first step to store STM in a silent state, and a second step to transform the silent state into an active state. This suggestion is in line with previous results that silent engrams have less spine density compared with active engrams^19,23^. However, further studies are needed to better understand the role of synaptic plasticity and to identify the downstream molecular players involved in this transformation.

Previous work has highlighted the necessity of post-learning sleep in memory consolidation, a process after which the engram can be actively and naturally recalled^28–30^. In line with this, we found that the silent STM trace is also stored after a period of post-learning sleep, whereby both generalized sleep deprivation and specified neuronal silencing during NREM disrupted the availability of this trace for future consolidation. Moreover, memory reactivation during sleep has been previously suggested as a mechanism underlying memory consolidation^31,33^. Here, we showed for the first time that the silent STM trace is reactivated during post-learning sleep, similar to its activated trace, which demonstrates that sleep stores both silent and active engrams in a similar manner. Furthermore, sleep deprivation has been shown to impair hippocampal protein synthesis and consequently memory consolidation^47^. This provides a direct connection between our finding that both post-learning sleep and post-learning protein synthesis are essential in the long-term storage of STM, with the former acting as a pre-requisite for the later. Taken together, our results shed light on the crucial role of sleep in the storage of a silent STM trace.

Finally, based on our results, we propose a model for STM consolidation in a two-step process (Extended Data Fig. 7). First, STM is stored as a silent engram through post-learning NMDAR activity, new protein synthesis, and offline NREM reactivations. Second, STM is consolidated as an active engram by re-exposure within a few days of the initial event through NMDAR activity and offline NREM reactivations.

## Methods

### Animals

Male c-fos::tTA/KA1::Cre double transgenic mice^38^ were used in the optogenetic recall experiment. Male C57BL/6J mice were used in the optogenetic inhibition experiment. Male KA1::Cre mice^38^ were used for other behavioral and pharmacological experiments. For *in vivo* calcium imaging, male C57BL/6J mice (n = 4) and male Thy1-GCaMP7 mice with C57BL/6J background (n = 1) were used^44^. Mice were maintained on a 12 h light-dark cycle at 24 ± 3°C and 55 ± 5% humidity with ad libitum access to food and water. All mice were aged 12–20 weeks at the time of behavioral experiments. All procedures involving the use of animals complied with the guidelines of the National Institutes of Health and were approved by the Animal Care and Use Committee of the University of Toyama, Toyama, Japan.

### The Novel Object Location task

The Novel Object Location (NOL) memory test was performed as described previously^23^ with some modifications.

#### Handling and Habituation

Mice were subjected to 6 minutes handling and 6 minutes habituation, during which they freely explored an empty arena without objects for 4 consecutive days.

#### Training

Mice were placed in the arena with two identical objects positioned in two different corners adjacent to the walls. Mice freely explored the objects for 15 minutes in the case of strong training or 6 minutes in the case of weak training. In the two-training protocol, mice were subjected to a second training session that was identical to the first training session in terms of context, object locations, and duration of exposure.

#### Test

The test was conducted either after 30 minutes for short-term retention or 24 hours for long-term retention. Mice explored the arena for 5 minutes, and one object had been moved to a different corner and the other object remained unmoved. A video tracking system (Muromachi Kikai, Tokyo, Japan) was used to record exploration behavior for later analysis.

#### Context and objects

The arena was a square; one wall was a transparent acrylic board covered with a white tape, and the three other walls were grey acrylic (width 29 cm, depth 25 cm, height 29 cm). The floor consisted of grey acrylic board covered with white hard plastic. The objects used were colored ceramic cups placed upside down (width 7.8 cm, depth 5.5 cm, height 12.5 cm). The position of the two objects was counterbalanced across mice within each group. The arena and the objects were thoroughly cleaned with 80% ethanol between mice.

#### Exploration analysis

The time spent exploring each object was manually counted and the exploration preference for each object was calculated as follows:

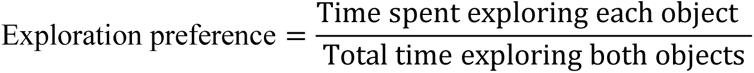

A greater exploration preference for the moved object than for the unmoved object was considered to indicate that mice had the ability to recall the object locations from the training session. Exploration preference was also calculated for the training sessions; mice that showed any bias towards either object (an exploration preference >0.6) were excluded from the dataset (n = 6). Exploration was defined as sniffing or touching objects with the nose. Touching by hands, jumping on, or moving around the objects were not considered as exploratory behaviors.

### Sleep detection

Electroencephalography (EEG) and electromyography (EMG) electrode placement and online sleep detection were performed as described previously^48^ with modifications.

#### Surgery

A custom-made 5-pin system that consisted of three wires terminating with screws was used; one screw was implanted in the parietal cortex for EEG recording, a ground screw and a reference screw were implanted in the cerebellar cortex, and two flexible wire cables were implanted in the neck muscle for EMG recording. All screws were fixed in place using dental cement. EEG and EMG signals were confirmed for each mouse individually before the start of experiments.

#### Data acquisition

All EEG/EMG recordings were performed using OpenEx Software Suite (Tucker Davis Technologies, USA) through a custom-made circuit file. EEG and EMG signals were amplified and filtered at 1–40 Hz for EEG and 65–150 Hz for EMG, and digitized at a sampling rate of 508.6 Hz.

#### Online sleep state differentiation

A custom-made circuit file was used to determine the sleep states by calculating the EMG root mean square (RMS) value, EEG delta power (1–4 Hz) RMS, EEG theta power (6–9 Hz), and RMS for 4 second epochs.

The criteria for defining a sleep state were as follows. Awake: an EMG RMS that was greater than the threshold value determined for each mouse. NREM: a delta/theta ratio that was greater than 1. REM: a delta/theta ratio that was less than 1.

Sleep state was defined if that state remained unchanged for three consecutive epochs (12 seconds). The sleep state defined by the program was confirmed by the experimenter through visual inspection of online waves and mouse activity, as follows.

Awake: mouse is moving or remains still + low amplitude and high frequency (8–30 Hz) EEG + high EMG activity. NREM: mouse remains still + high amplitude and low frequency (0.5–4 Hz) EEG + low EMG activity. REM: mouse remains still + low amplitude and high frequency (4–9 Hz) EEG + flat EMG activity; a tremor may occasionally appear.

#### Sleep deprivation

Sleep deprivation was conducted as previously described^47^ for the minimum induction of stress. Immediately after training, mice were returned to their home cages and the experimenter observed them for 5 consecutive hours. Whenever mice began to stand still or sleep, the experimenter gently tapped or pushed the home cage which alerted the mice and stopped them from sleeping.

### Pharmacological experiments

#### Drugs

Anisomycin (Sigma A9789, 62.5 μg/μl), a protein synthesis inhibitor, was dissolved with 1N HCl and adjusted to pH 7.4 with NaOH^37^. D-AP5 (Tocris Bioscience, 0106, 30 mM), an N-methyl-D-aspartate receptor blocker, was dissolved in phosphate buffered saline (PBS) (T900, Takara BIO Inc, Japan)^13^. Drugs were aliquoted into single-use tubes and stored at −20°C until use.

#### Stereotactic surgery

Mice (12 weeks) were anesthetized with a combination of three drugs (0.75 mg/kg, I.P., medetomidine (Domitor; Nippon Zenyaku Kogyo Co., Ltd., Japan), 4.0 mg/kg, I.P., midazolam (Fuji Pharma Co., Ltd., Japan), and 5.0 mg/kg, I.P., butorphanol (Vetorphale, Meiji Seika Pharma Co., Ltd., Japan))^49^. Mice were then placed on a stereotactic apparatus (Narishige, Japan), guide cannulas (C313GS-5/SPC, gauge 22, Plastics One, USA) were implanted bilaterally into the hippocampal CA1 region (AP -2.0 mm, ML ±1.4 mm, DV 1.0 mm from bregma), and then dummy cannulas (C313IDCS-5/SPC, zero projection, Plastics One, USA) were inserted into the guide cannulas to protect them from dust. Micro screws were fixed near the bregma and lambda, and the guide cannulas were fixed in position using dental cement. Mice had a recovery period of 4 weeks in individual home cages before the start of the behavioral experiments.

#### Micro-infusion after training session

Immediately after training, mice were anesthetized with isoflurane (099-06571, FUJIFILM Wako Pure Chemical, Osaka, Japan), and injection cannulas (C313IS-5/SPC, Plastics One, USA) were placed into guide cannulas projecting 0.5 mm below the tip of the guide cannulas. Mice were then returned back to their home cages. The internal cannulas were attached through a thin plastic tube filled with water to 10 μl Hamilton syringes that were fixed to an automated pump to maintain the drug flow rate at 0.2 μl/min. A total volume of 1 μl (anisomycin or D-AP5 as drugs, and PBS as a control) was injected bilaterally into the CA1 region, and the injection cannulas were left in place for 5 minutes after infusion to allow for drug diffusion. Mice were perfused 1 day after the test and the brain was extracted for histological examination.

### Optogenetic experiments

#### Viral vectors

For the optogenetic recall experiment, AAV_9_-TRE2G::DIO-ChR2(T159C)-mCherry (Titer: 1.3 × 10^13^ vg/ml)^38^ was used. For the optogenetic inhibition experiment, AAV_9_-CaMKII-ArchT-eYFP (Titer: 3.02 × 10^16^ vg/ml, a gift from Dr. R. Okubo-Suzuki) was used. The recombinant AAV_9_ production was performed using a minimal purification method, and viral genomic titer was subsequently calculated as described previously^50^. Briefly, pAAV recombinant vector was produced using HEK293 T-cells (AAV293; 240073, Agilent Tech, CA, USA) cultured in 15 cm dishes (Corning, NY, USA). Cultured cells were maintained in Dulbecco’s Modified Eagle Medium (D-MEM) (11995-065, GIBCO Life Technologies, USA) supplemented with 10% fetal bovine serum (FBS) (10270106, GIBCO Life Technologies, USA), 1% 2 mM L-Glutamine (25030-149, GIBCO Life Technologies, USA), 1% 10 mM non-essential amino acid (MEM NEAA 100x, 11140-050, GIBCO Life Technologies, USA), and 1% (100x) penicillin-streptomycin solution (15140-148, GIBCO Life Technologies, USA). Confluent (70%) HEK293 T-cells were transfected using medium containing the constructed expression vector, pRep/Cap, and pHelper (240071, Agilent Technologies, Santa Clara, CA, USA) mixed with the transfection reagent polyethylenimine hydrochloride (PEI Max, 24765-1, Polysciences, Inc., Warrington, PA, USA) at a 1:2 ratio (W/V). After 24 h, the transfection medium was discarded, and cells were incubated for another 5 days in an FBS-free maintenance medium. On day 6, the AAV-containing medium was collected and purified from cell debris using a 0.45 μm Millex-HV syringe filter (SLHV033RS, Merck Millipore, Germany). The filtered medium was concentrated and diluted with D-PBS (14190-144, GIBCO Life Technologies, USA) twice using the Vivaspin 20 column (VS2041, Sartorius, Germany) after blocking the column membrane with 1% bovine serum albumin (01862-87, Nacalai Tesque, Inc., Japan) in PBS. To further calculate the titer, degradation of any residual cDNA in the viral solution from production was first assured by benzonase nuclease treatment (70746, Merck Millipore, Germany). Subsequently, viral genomic DNA was obtained after digestion with proteinase K (162-22751, FUJIFILM Wako Pure Chemical, Osaka, Japan), extraction with phenol/chloroform/isoamyl alcohol 25:24:1 v/v, and then precipitation with isopropanol and final dissolution in TE buffer (10 mM Tris [pH 8.0], 1 mM EDTA). Titer quantification for each viral solution, referenced to that of the corresponding expression plasmid, was done using real-time quantitative PCR using THUNDERBIRD SYBR qPCR Master Mix (QRS-201, Toyobo Co., Ltd, Japan) with the primers 5′-GGAACCCCTAGTGATGGAGTT-3′ and 5′-CGGCCTCAGTGAGCGA-3′ targeting the inverted terminal repeat (ITR) sequence. The cycling parameters were adjusted as follows: initial denaturation at 95°C for 60 sec, followed by 40 cycles at 95°C for 15 sec and at 60°C for 30 sec. For the control experiment, AAV_1_-CaMKII-eYFP (titer 1.9 × 1013 vg/ml, Addgene, #105622) was used.

#### Stereotactic surgery

Mice (12 weeks) were anesthetized and placed on a stereotactic apparatus as mentioned in the pharmacological experiments AAV (0.5 μl) was injected bilaterally into the hippocampal CA3 region (AP -2.0 mm, ML ±2.3 mm from bregma, DV 2.0 mm from the dura) or the hippocampal CA1 region (AP -2.0 mm, ML ±1.4 mm, DV 1.5 mm from bregma) using a glass micropipette filled with mineral oil attached to a 10 μl Hamilton syringe. The flow rate was fixed at 0.1 μl/min using a microsyringe pump and its automated controller (Narishige, Tokyo, Japan). The glass micropipette was left in place for 5 minutes after virus injection. Guide cannulas (C316GS-5/SPC, gauge 24, Plastics One, USA) were implanted bilaterally into the hippocampal CA1 region (AP -2.0 mm, ML ±1.4 mm, DV 0.5 mm from bregma) and were covered with dummy cannulas (C316IDCS-5/SPC, zero projection, Plastics One, USA) to protect them from dust. Micro screws were fixed near the bregma and lambda, and the guide cannulas inserted were fixed using dental cement. Mice were maintained in individual home cages on 40 mg/kg Dox (doxycycline) for c-fos::tTA/KA1::Cre double transgenic mice or normal food pellets for C57BL/6J mice, and allowed to recover for 4–5 weeks before the start of behavioral experiments.

#### Photo-stimulation during the test

Mice were maintained on Dox during the handling and habituation sessions, and then were taken off Dox 48 hours before the training session and remained off Dox until sacrificed. On the test day, mice were anesthetized with isoflurane, and internal cannulas were replaced by a two-branch optical fiber unit consisting of a plastic cannula body and a 0.25 mm diameter optic fiber (COME2-DF2-250; Lucir, Ibaraki, Japan), which were placed inside the guide cannulas such that the tip of the optical fiber was targeted slightly above the CA1 region (DV 1.0 mm from bregma). Mice were returned to their home cages for 1 hour to recover from the anesthesia, and then moved to the experimental room where the fiber unit was fixed to an optical swivel above the test context (COME2-UFC; Lucir) that was connected to a laser source (200 mW, 473 nm, COME-LB473/200; Lucir). Pulses were delivered during the test session (5 minutes) according to a pre-fixed schedule using a stimulator (COME2-SPG-2; Lucir) in a time-lapse mode. Mice received 20 pulses of 473 nm blue light every second (20 Hz); each pulse had a duration of 0.5 msec and an inter-pulse interval of 49.5 msec. Mice were perfused 90 minutes after the test and their brains were extracted for histological examination.

#### Photo-inhibition during sleep

Immediately after the first training, mice were anesthetized with isoflurane and optic fibers were attached, as explained above, and then an EEG/EMG recording unit was attached. Mice were then placed into their home cages to sleep. For a duration of 2 hours from sleep onset, 589 nm orange light was delivered above the hippocampal CA1 region, as explained above, whenever the mouse entered the NREM sleep stage. Mice were perfused 1 day after the test and their brains were extracted for histological examination.

### *In vivo* calcium imaging experiment

#### Viral vector

AAV_9_-CaMKII::G-CaMP7 (titer: 9.4 × 10^12^ vg/mL)^49^ after 40-fold dilution with PBS was used.

#### Stereotactic operation

All surgical steps were performed as previously described^44,49,51,52^. Mice (12 weeks) were anesthetized and the AAV (0.5 μl) was injected unilaterally into the right hippocampal CA1 region (AP -2.0 mm, ML 1.4 mm, DV 1.5 mm from bregma), as described above, and returned to their home cages (this step was skipped when Thy1-GCaMP7 transgenic mice were used). One week after AAV injection, mice were re-anesthetized and placed again on the stereotactic apparatus, and a craniotomy (2.0 mm diameter) was drilled over the center of the injection site. The neocortex and corpus callosum above the alveus overlying the dorsal hippocampal CA1 region were aspirated under constant irrigation with saline using a 26-gauge flat-tip needle. Saline was applied to control the bleeding. A gradient index (GRIN) lens (1 mm diameter, 4 mm length; Inscopix, CA, USA) was placed on the center of the alveus using handmade forceps attached to a manipulator (Narishige, Tokyo, Japan). Bone wax was melted by a low-temperature cautery and applied to seal any gaps between the skull edge and the GRIN lens, and the lens was fixed in place using dental cement. The EEG/EMG custom-made 5-pin system was attached and cemented into the skull as mentioned earlier. An additional four anchor screws were planted on the skull for further fixation. A plastic cover was fixed around the GRIN lens for protection until the next surgery, and mice were returned to their home cages. Three weeks after GRIN lens implantation, mice were re-anesthetized and placed back onto the stereotactic apparatus to set a baseplate (Inscopix, CA, USA). A gripper (Inscopix, CA, USA) holding the baseplate attached to a miniature microscope (nVista HD v3; Inscopix, CA, USA) was lowered over the previously set GRIN lens until visualization of clear blood vessels was possible, which indicated the optimal focal plane. Dental cement was then applied to fix the baseplate in this position. Mice were returned to their home cages and allowed a recovery period of 1 week before the start of behavioral experiments.

#### Calcium imaging during the NOL task

Handling was done for 6 minutes, as mentioned above, then the miniature microscope was mounted onto the baseplate along with the EEG/EMG recording unit. Mice were left in their home cages for 10 minutes before the habituation session started.

For the first 3 days of habituation, no calcium imaging was performed; however, the miniature microscope was attached so that mice could habituate to the context exploration with the mounted microscope. Imaging was performed during the last habituation session (day 4) for 6 minutes, during the first and second training sessions (6 minutes each), and during the test session (6 minutes: n = 3; 5 minutes: n = 2). For imaging during post-learning sleep after the first and second training sessions, sleep state was determined using EEG/EMG monitoring, as explained above, and imaging was performed for the first 5 minutes of NREM sleep.

#### Data acquisition, processing, and cell detection

Calcium transients were captured at 20 Hz using nVista acquisition software (Inscopix, CA, USA) at the maximum gain and optimal LED power (the value was adjusted for each mouse). Movie recordings of all sessions were extracted from nVista Data acquisition (DAQ) box (Inscopix, CA, USA) and temporally stitched together (concatenated) to create one movie with all recorded sessions combined one after the other using Inscopix data processing software (Inscopix, CA, USA). Using the same software, the movie was spatially down sampled by a factor of 2, and then motion correction was applied to remove motion artifacts. A reference frame with clear blood vessels as the landmark was chosen, and all other frames were aligned to it. Then, as previously described^44,49^, using Mosaic software (Inscopix, CA, USA), additional motion correction was applied to the movie using the same reference frame. Using Fiji software (NIH), the movie was temporally divided into individual sessions and low bandpass filtered to reduce noise. Using Mosaic software (Inscopix, CA, USA), the fluorescence signal intensity change (ΔF/F) for each session was calculated according to the formula ΔF/F = (F – Fm)/Fm, where F represents each frame’s fluorescence and Fm is the mean fluorescence for the whole session. Movies representing each session were re-concatenated again to generate one movie for all sessions in the ΔF/F format. Finally, cells were identified using an automatic sorting system, HOTARU^45^, and each cell’s calcium signals over time were extracted in a (Ď; time × neuron) matrix format.

#### Data analysis

Data analysis was performed using a custom-made MATLAB code. Calcium signals were subjected to high-pass filtering (0.01 Hz threshold) to remove low frequency fluctuations and background noise. Mean z-scores were calculated for each session using a cut-off value of <3 SD from the local maxima of the ΔF/F signal of each session. The mean z-score of each session was normalized to that of the habituation session, and training-1-emerging cells were sorted (cells that were not activated during the habituation session, but started to become active during training-1; a cell was considered as a training-1-emerging cell if it showed a 2-fold higher response to the training-1 session than to the habituation session). Activity of training-1-emerging cells were tracked across the following sessions.

To measure the reactivation of training cells in post-learning NREM sleep, the percentage of overlapping cells was calculated and compared to chance levels as follows:

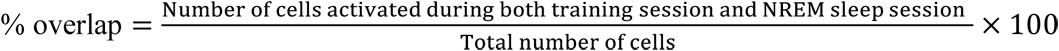

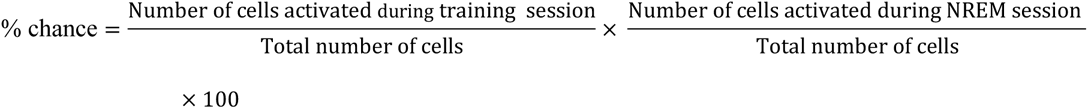

### Histology

#### For virus-infected animals

Mice were deeply anesthetized with an overdose of pentobarbital solution and transcardially perfused using 4% paraformaldehyde in PBS, pH 7.4. The brains were then removed and post-fixed by immersing in 4% paraformaldehyde in PBS for 24 hours at 4°C, equilibrated in 25% sucrose in PBS for 2 days, and then frozen in dry-ice powder and stored at -80°C until sectioning. Coronal sections of a 50 μm thickness were cut using a cryostat and then transferred to 12-well cell culture plates with 5 sections/well (Corning, Corning, NY) containing PBS solution. The floating sections were then treated with 4,6-diamidino-2-phenylindole (DAPI, 1 μg/ml, 10236276001; Roche Diagnostics) at room temperature for 20 minutes, and then washed 3 times with PBS (3 minute/wash). The DAPI-stained sections were then mounted on a glass slide with ProLong Gold antifade reagent (Invitrogen, Thermo Fisher Scientific, Waltham, MA).

Images of native mCherry, eYFP, and GCaMP fluorescence were acquired using a Zeiss LSM 780 confocal microscope with a Plan-Apochromat 20x or 5x objective lens (Nikon, Japan). All acquisition parameters were kept constant within each magnification and for all images. Unstained coronal sections were stored at –20°C in a cryoprotectant solution (25% glycerol, 30% ethylene glycol, 45% PBS) for further use if needed.

#### For drug-injected animals

Mice were sacrificed by decapitation, and brains were rapidly extracted and immediately frozen in dry-ice powder and stored at –80°C until sectioning. Coronal sections of 30 μm thickness were cut using a cryostat, mounted onto glass slides and air dried, and then examined to confirm the injection trace in the hippocampal tissue. Mice were excluded if the injection trace was not clear (n = 4).

### Statistical analysis

Statistical analysis was performed using GraphPad Prism 6 (GraphPad Software, Inc., USA). Comparisons of data between two groups were performed using Student’s t-test (two-tailed) and the Wilcoxon rank sum test. Multiple-group comparisons were assessed using a two-way analysis of variance (ANOVA) followed by post-hoc Bonferroni multiple comparisons test. Quantitative data are expressed as the mean ± SEM, except for Fig. 6d, right; data are presented using box plots.

### Data and code availability

The data and codes that supported the findings of this study are available from the corresponding author upon reasonable request.

## Acknowledgements

We thank K. Deisseroth (Stanford University) for the ChR2 (T159C) AAV vector and the ArchT-eYFP plasmid, A. Konno and H. Hirai for disclosing the AAV_9_ virus production protocol prior to publishing, and R. Okubo-Suzuki for preparation of the AAV_9_-CaMKII-ArchT-eYFP viral vector. We thank M.H. Aly for technical assistance, S. Tsujimura and S. Okami for maintaining mice, K. Choko and Y. Saitoh for preparation of EEG/EMG electrodes, N. Oishi for technical teaching, and all members of the Inokuchi Laboratory for discussions and suggestions. This work was supported by JSPS KAKENHI (JP18H05213), the Core Research for Evolutional Science and Technology (CREST) program (JPMJCR13W1) of the Japan Science and Technology Agency (JST), a Grant-in-Aid for Scientific Research on Innovative Areas “Memory dynamism” (JP25115002) from MEXT, and the Takeda Science Foundation (to K.I.). Additional support was provided by JSPS KAKENHI (20H03554 and 17K19445), THE HOKURIKU BANK Grant-in-Aid for young scientists, THE FIRSTBANK OF TOYAMA Scholarship Foundation research grant, and the Takeda Science Foundation (to M.N.). Grant-in-Aid for young scientists from JSPS KAKENHI (JP 19K16892) (to K.A.). The Otsuka Toshimi Scholarship Foundation supported M.W.

## Author contribution

M.W., M.N., K.A., and K.I. designed the experiments and wrote the manuscript. M.W. performed all experiments. M.W. and M.N. analyzed the data. M.N. generated MATLAB codes. K.I. supervised the entire project.

## Competing interests

The authors declare no competing interests.

**Correspondence and requests for materials** should be addressed to K.I.

---

**Extended Data Fig. 1.**
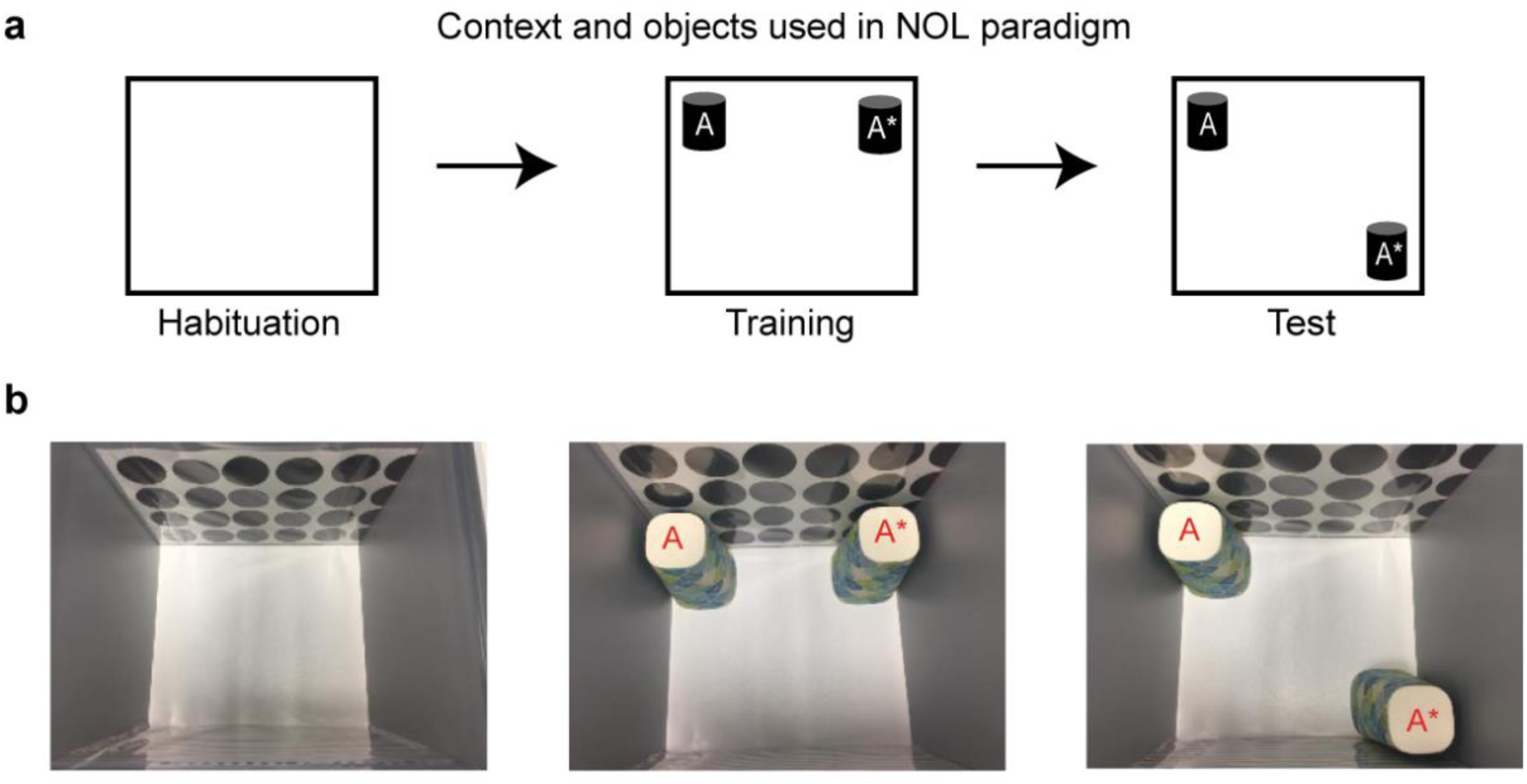
Novel object location paradigm layout. **a,** Context and objects layout; object A: remains in its location during test, object A*: changes its location during test. Object location was counterbalanced across mice. **b,** Photos of context and objects used, 2 walls of the context were covered with spatial cues to aid the mice in navigation, one wall had circular patterns and the opposite wall had stripes pattern. (see Methods for a full description of the NOL paradigm).

**Extended Data Fig. 2.**
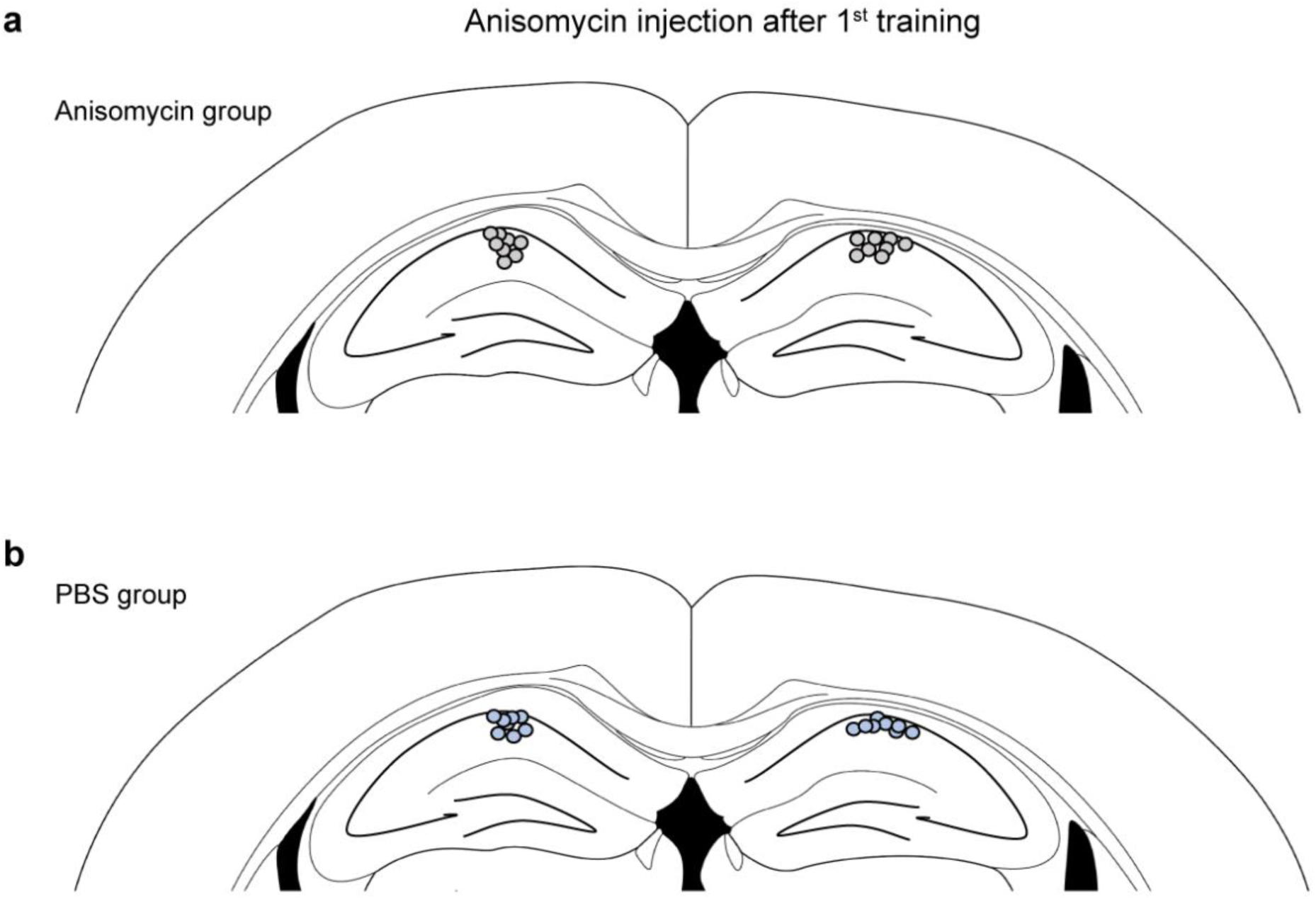
Cannula tip placement in mice infused with anisomycin and PBS after the first training. **a,** Anisomycin group injection traces (n = 8). **b,** PBS group injection traces (n = 8).

**Extended Data Fig. 3.**
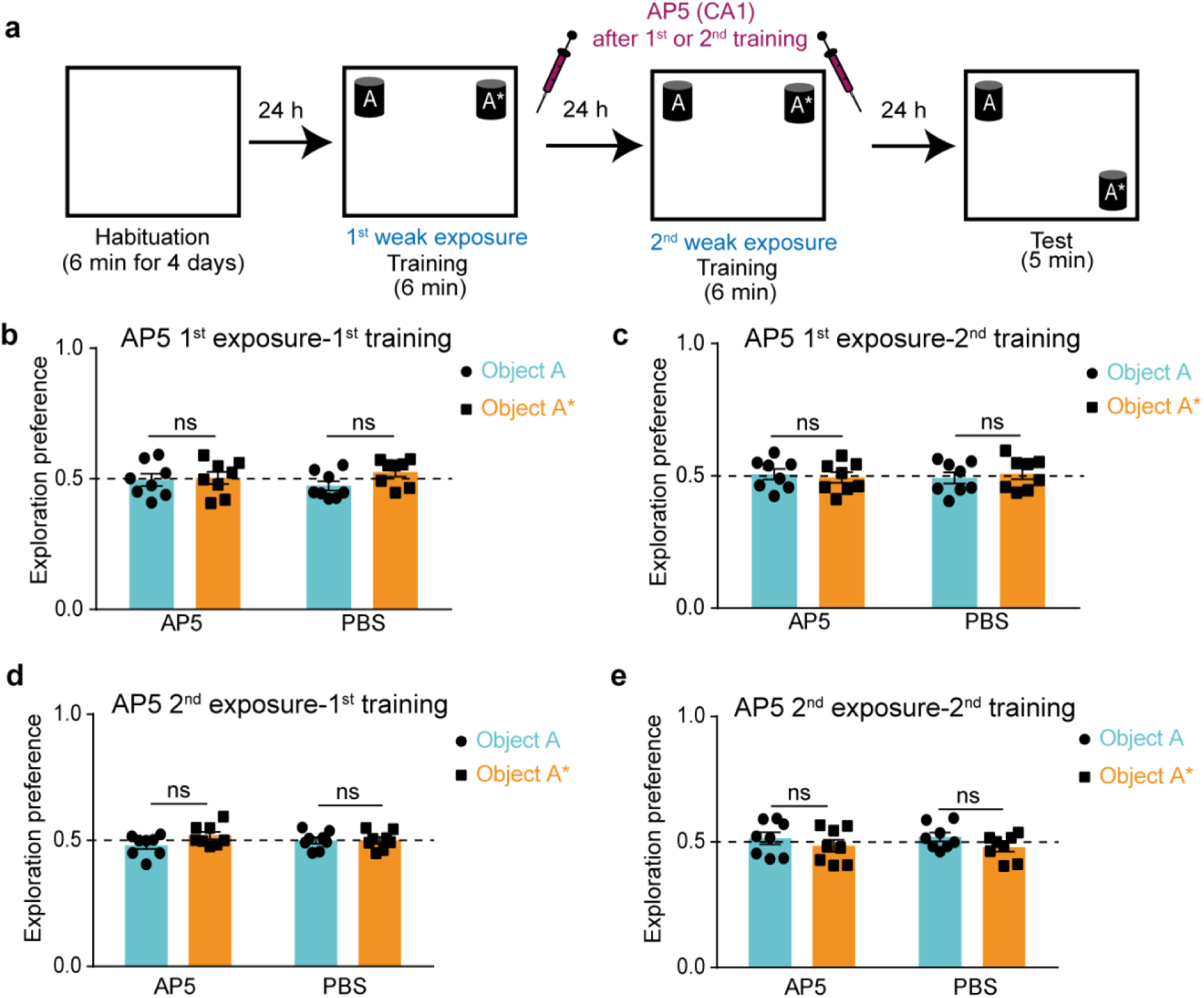
Training sessions for N-methyl-D-aspartate receptor blockade experiments. **a,** NOL two-training paradigm with 1-day interval in between with AP5 injection after either the first or the second training sessions into the CA1 region. **b and c,** Exploration preference for each object during the first **(b)** or the second **(c)** training sessions for AP5- (n = 8) and PBS- (n = 8) injected groups after the first training session. **d and e,** Exploration preference for each object during the first **(d)** or the second **(e)** training sessions for AP5- (n = 8) and PBS- (n = 8) injected groups after the second training session. Comparisons were made using unpaired student’s t-test; ns, not significant (*P*> 0.05). Data are presented as the mean ± SEM.

**Extended Data Fig. 4.**
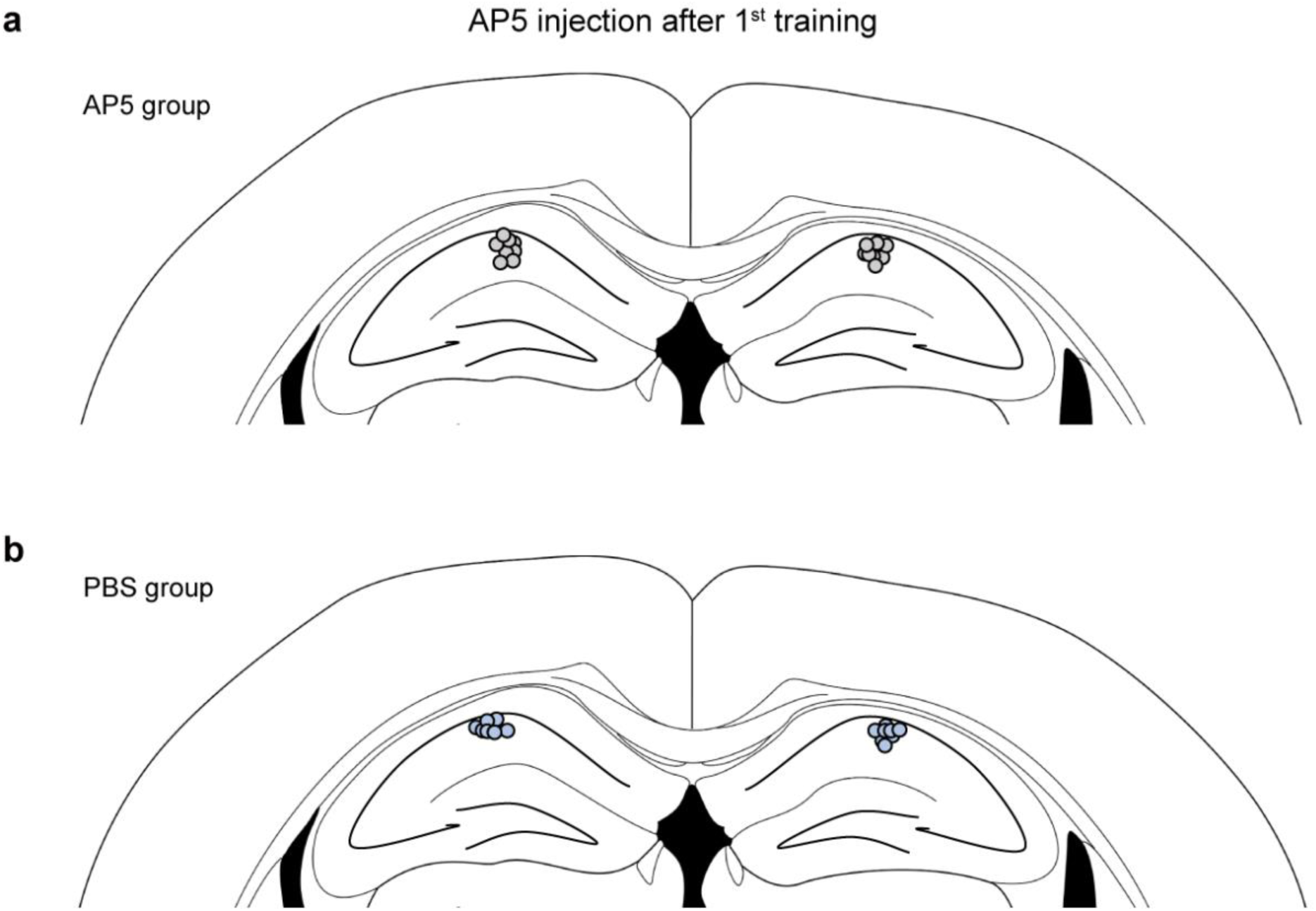
Cannula tip placement in mice infused with AP5 and PBS after the first training. **a,** AP5 group injection traces (n = 8). **b,** PBS group injection traces (n = 8).

**Extended Data Fig. 5.**
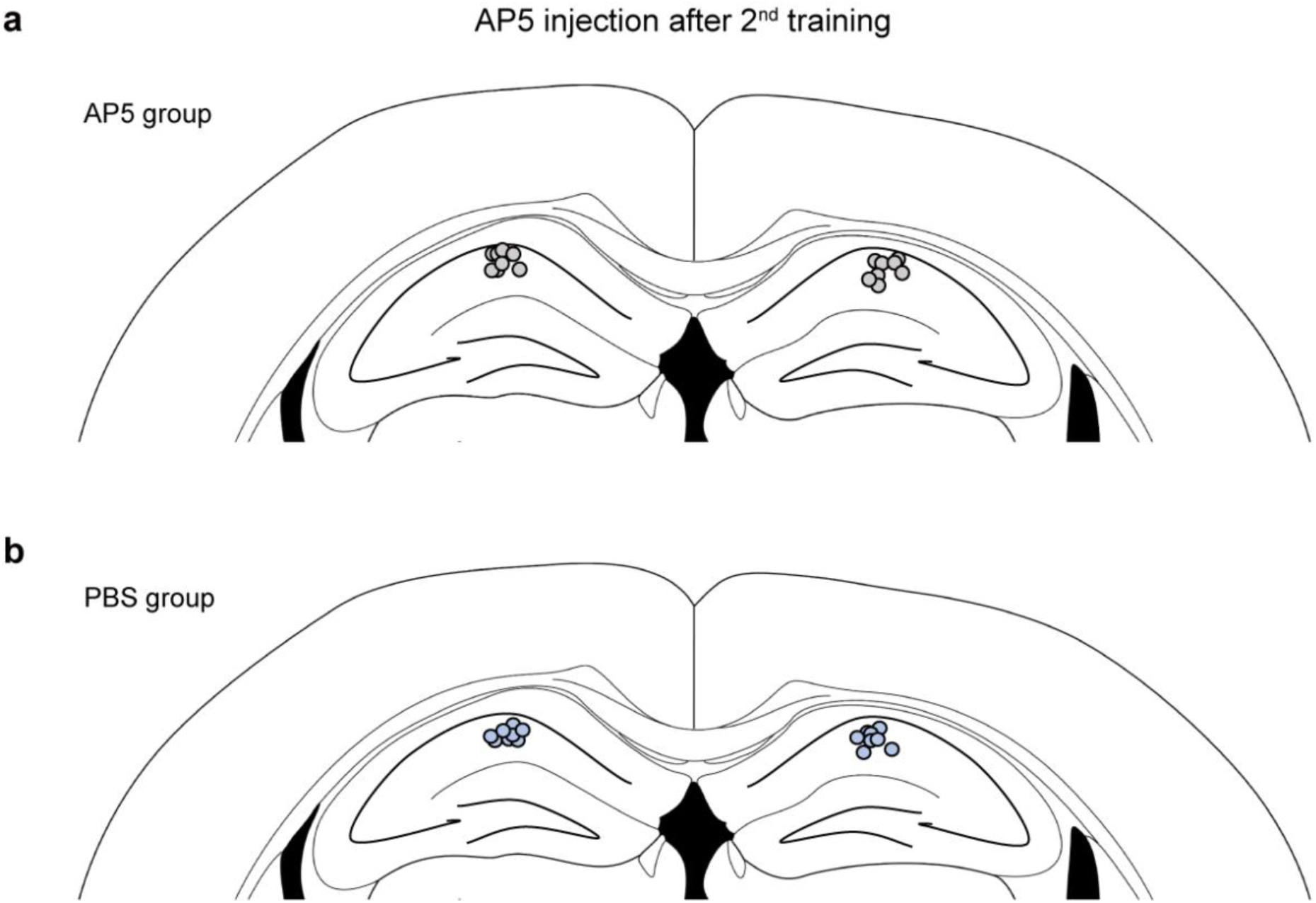
Cannula tip placement in mice infused with AP5 and PBS after the second training. **a,** AP5 group injection traces (n = 8). **b,** PBS group injection traces (n = 8).

**Extended Data Fig. 6.**
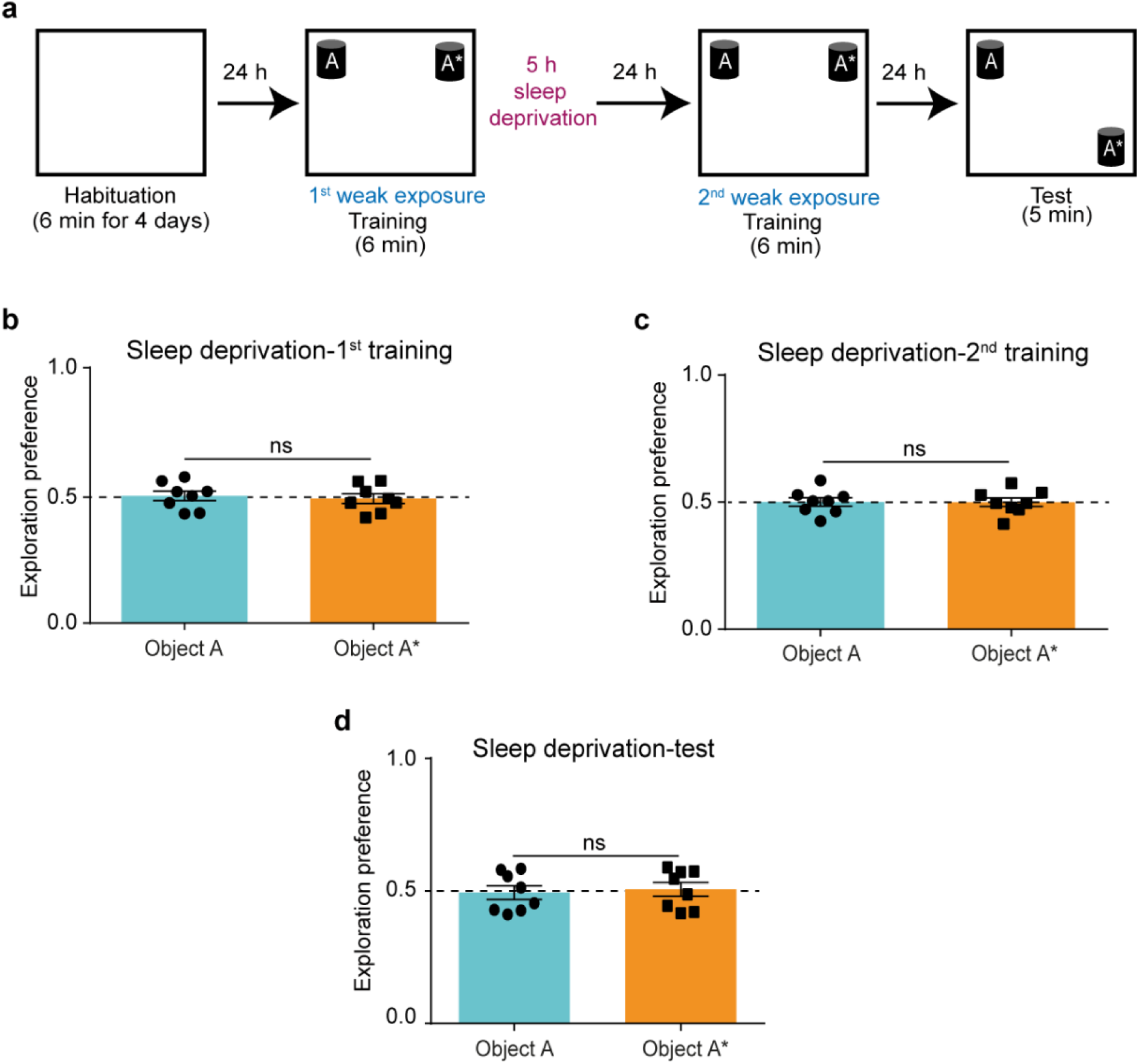
Post-learning sleep is required to preserve STM trace. **a,** NOL two-training paradigm with 1-day interval with 5-hour sleep deprivation after the first training session. **b and c,** Exploration preference for each object during the first **(b)** and the second **(c)** training sessions (n = 8). **d,** Exploration preference for each object during the test session (n = 8). Comparisons were made using unpaired student’s t-test; ns, not significant (*P*> 0.05). Data are presented as the mean ± SEM.

**Extended Data Fig. 7.**
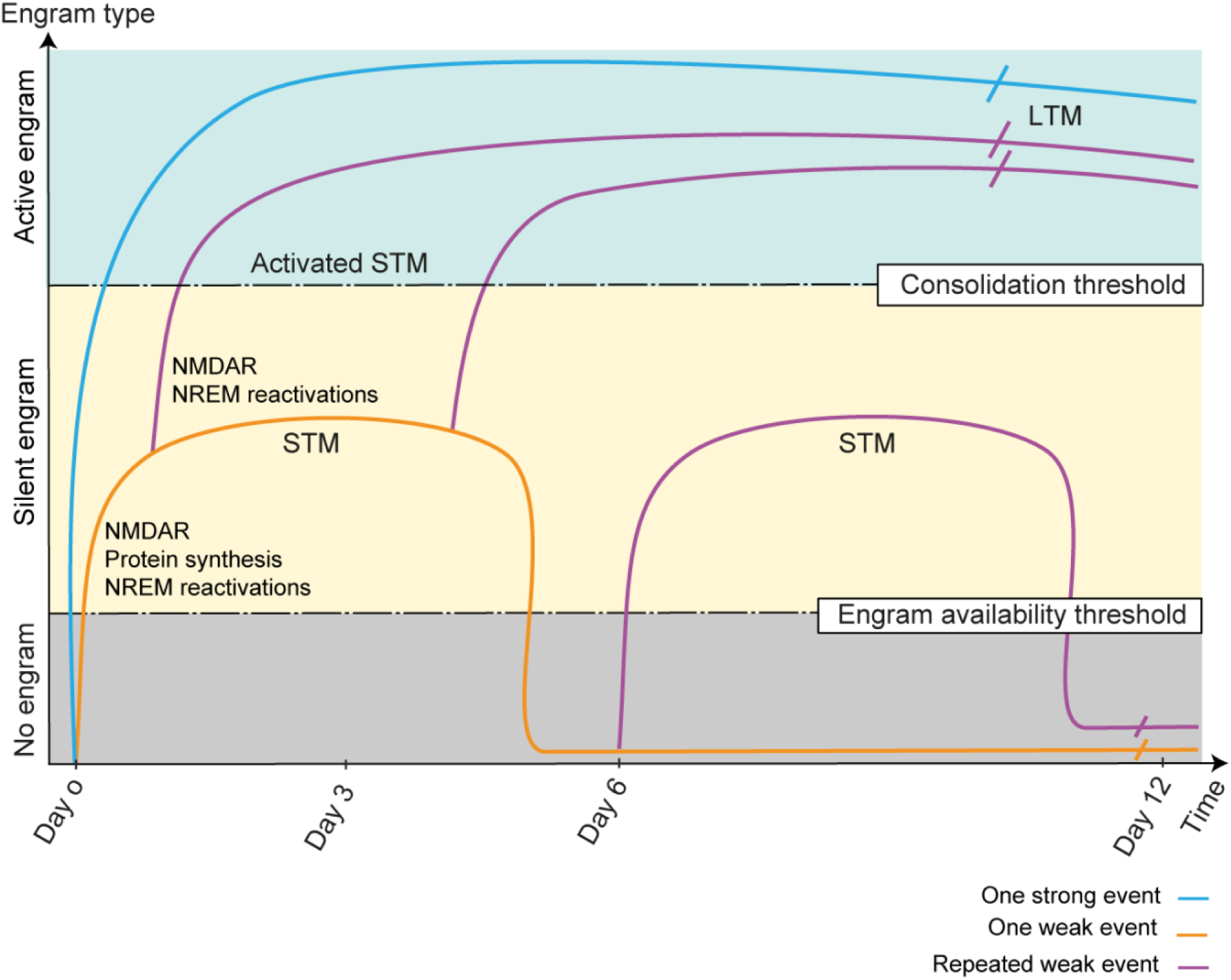
Hypothesized model for STM vs. LTM engrams. One strong event is sufficient to consolidate an LTM, which can be actively recalled through its active engram. One weak event is not sufficient for consolidation; however, it forms an STM stored in the form of a silent engram through post-learning NMDAR activation, new protein synthesis, and NREM sleep reactivations. A repeated weak event then activates this silent engram by consolidation within its lifetime (<6 days) through post-learning NMDAR activation and NREM sleep reactivations. However, if the second weak event is repeated after 6 days, the first STM engram is no longer available and is then processed as a new event by forming a new silent STM engram with similar properties.

## Notes

### Competing Interest Statement

The authors have declared no competing interest.

